# STING: A Graph Neural Network Approach for Computational Inference of Spatial Transcriptomic Profiles

**DOI:** 10.1101/2025.04.23.650236

**Authors:** Kaushik Karambelkar, Arvind Rao, Mayank Baranwal

## Abstract

Spatial transcriptomics enables the measurement of mRNA counts at spatial locations within a tissue but faces challenges such as high experimental costs, technical expertise requirements, and low RNA detection efficiency at high resolution. We present **STING** (Spatial Transcriptomics Inference using Graph neural networks), a computational approach that infers spatial gene expression patterns with high efficiency. STING integrates convolutional neural networks (CNNs) pre-trained on histological images with graph neural networks (GNNs) to model spatial proximity, representing tissue sections as nearest-neighbor graphs where spatially adjacent spots are interconnected. We train GNNs on these graphs to predict gene expression at additional tissue locations, including unseen samples. Evaluated on two public spatial transcriptomics datasets (59 and 36 tissues), STING achieves a Pearson correlation coefficient (PCC) of up to **0.79** for super-resolution and **0.69** for tissue-wide inference, outperforming existing methods in accuracy. Our results demonstrate that STING is an effective and computationally efficient tool for predicting spatial gene expression, with significant applications in cancer research, personalized medicine, and beyond.

## Introduction

The human body comprises over a hundred distinct cell types, all sharing the same genome^1^. Understanding cellular diversity and classification is crucial for deciphering tissue function and disease mechanisms^2^. While proteins best define cell identity and function, large-scale protein profiling remains impractical^3,4^. Instead, gene expression analysis serves as a feasible alternative, following the central dogma of molecular biology.

Bulk RNA sequencing is well-established in molecular biology, with applications spanning disease characterization and pharmacogenomics^5,6^. However, it provides only aggregated gene expression signals, missing critical cell-type-specific details^7,8^. Single-cell RNA sequencing (scRNA-seq), introduced in 2009^9^, allows transcriptomic profiling at the cellular level but requires isolating viable single cells, which is labor-intensive and impractical for certain cell types, such as neurons^10,11^. Moreover, scRNA-seq disrupts the native tissue context, where intercellular interactions influence gene expression^1^.

Spatially resolved transcriptomics (SRT)^12,13^ overcomes these limitations by capturing spatial gene expression patterns across tissues and was named “Method of the Year” in 2020 by *Nature Methods*^14^. This can be achieved through probe-based sequencing or imaging techniques like in-situ sequencing (ISS) and in-situ hybridization (ISH)^15^. SRT provides a comprehensive view of cellular interactions within tissues, with significant applications in cancer research, particularly for studying tumor microenvironments and intra-tumor heterogeneity^16,17^. However, high-resolution SRT is costly and constrained by trade-offs between spatial resolution, gene coverage, and RNA detection efficiency^1,13^.

Whole-slide images (WSIs) of H&E-stained tissues offer an alternative, capturing cellular organization and morphological features in high resolution^18,19^. Machine learning (ML) has proven effective for WSI analysis in cancer research, supporting tasks such as tumor classification^20,21^, cancer subtyping^22,23^, and mutation prediction^24,25^. With the advent of SRT, ML-driven approaches have emerged for inferring spatial gene expression from WSIs^26–28^, offering a more accessible alternative to direct spatial transcriptomic profiling. Typically, these methods align WSIs with tissue spots where gene expression is measured and predict gene expression patterns using local image features.

Convolutional neural networks (CNNs) have been widely used for spatial gene expression inference, leveraging pre-trained architectures like VGG-16^29^ and `DenseNet-121`^30,31^. However, CNNs are not adequate at incorporating spatial proximity, treating all image regions independently despite the fact that adjacent spots often exhibit similar gene expression patterns.

Moreover, CNNs pre-trained on natural images may not optimally capture features from H&E-stained tissues, and training them from scratch requires extensive data and computational resources.

Alternative ML models incorporating vision transformers, convolutional autoencoders, and graph neural networks (GNNs) have been explored^28,32–35^. While these methods attempt to leverage spatial relationships in gene expression, they often suffer from high computational costs or lack interpretability in a biological context. This underscores the need for an efficient and biologically relevant approach to spatial transcriptomic inference.

In this paper, we introduce **STING** (*Spatial Transcriptomics INference using Graph neural networks*), a simple yet powerful two-component framework for inferring spatial gene expression from H&E-stained tissue images. STING harnesses the feature extraction capabilities of convolutional neural networks (CNNs) pre-trained specifically on histological images and integrates graph neural networks (GNNs) to capture spatial dependencies in gene expression across tissue sections. We demonstrate STING’s ability to generate high-resolution spatial gene expression maps from lower-resolution counterparts and infer transcriptomic profiles in previously unseen tissue sections. Across two publicly available spatial transcriptomics datasets, STING consistently outperforms existing methods in most tissue sections. Moreover, STING is computationally efficient, requiring fewer trainable parameters than alternative approaches while maintaining high interpretability.

Our key contributions can be summarized as follows:

- We develop a **graph neural network model** that infers spatial gene expression from histopathological images without the need for extensive computational resources.
- Our model effectively **super-resolves spatial gene expression**, enhancing resolution from lower-resolution expression maps using tissue images.
- STING is capable of **predicting spatial transcriptomic profiles** for tissues where no prior gene expression data is available.

By bridging histological imaging and spatial transcriptomics, STING provides an accessible and efficient solution for high-resolution gene expression inference, with broad applications in biomedical research and precision medicine.

## Results

### Datasets

We use two publicly available spatial transcriptomics datasets for training and validation. The first dataset, **Her2-ST**, from Andersson et al.^36^, includes WSIs of 36 tissue sections from eight Her2-positive breast cancer patients. The second dataset, **ST-Net**, from He et al.^26^, consists of spatial transcriptomics data and corresponding WSIs for 68 tissues from 23 mixed-subtype breast cancer patients. Spatial gene expression profiles in both datasets are generated using the 10x Visium platform.

Each tissue is represented by its H&E-stained image, pixel coordinates of the spots where gene expression is measured, and corresponding gene expression scores. Each spot is identified by its position within the 10x Visium grid, allowing precise mapping of spatial coordinates to gene expression values. The Her2-ST dataset includes gene expression scores for **14,800–15,900** genes, while the `ST-Net` dataset covers **16,700–19,800** genes.

Since these datasets are publicly available, no ethical approvals are required. The sources of all datasets used are detailed in the Data & Code Availability section.

### Super-Resolution of Gene Expression

We super-resolve gene expression for 250 genes in one tissue section selected from each patient in both datasets. In the Her2-ST dataset, the highest-quality super-resolution is observed in tissue section *B1* (mean PCC = 0.74), while the lowest is in *F1* (mean PCC = 0.24). In the `ST-Net` dataset, the most optimal super-resolution occurs in tissue sections *BC23287_C1* and *BC23944_D2* (mean PCC = 0.47 for both), whereas the lowest is in *BC23277_D2* (mean PCC = 0.11). Figure 2a visualizes the PCCs for all tissues across both datasets.

**Figure 1.**
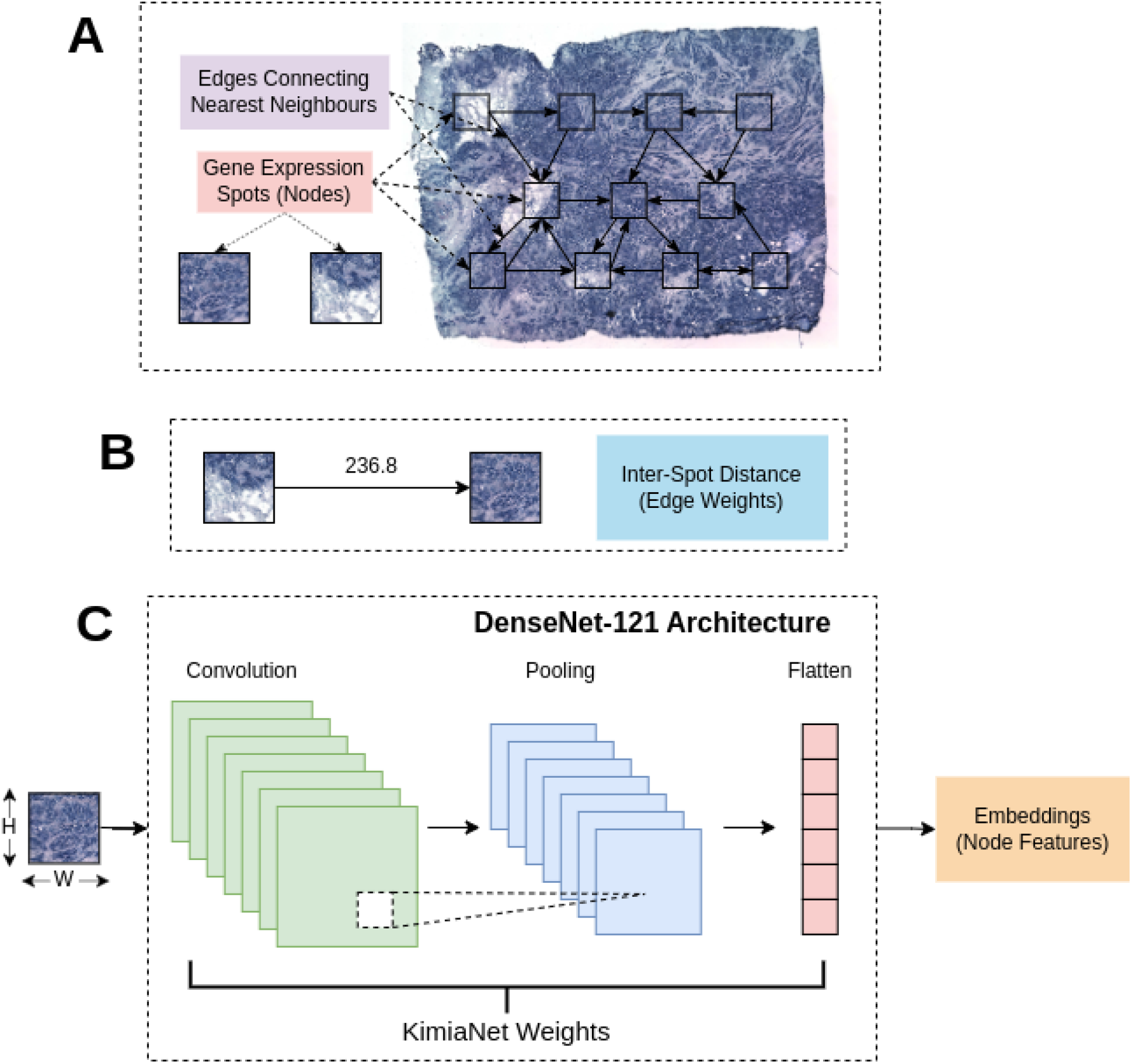
The process of generating graph data from a tissue. (A) Spots where gene expression is measured form the nodes in the graph, and each spot is connected via an edge to “k” of its nearest neighbors. (B) The weight of each edge is determined by the distance, in pixels, between the spots that it connects. (C) To generate node features, we removed the fully connected layer in `DenseNet-121`, and applied to it the weights of `KimiaNet`, a `DenseNet-121`architecture pre-trained on WSIs. An area around each spot, measuring WxH pixels was then passed through it to generate embeddings, which were used as node features.

**Figure 2.**
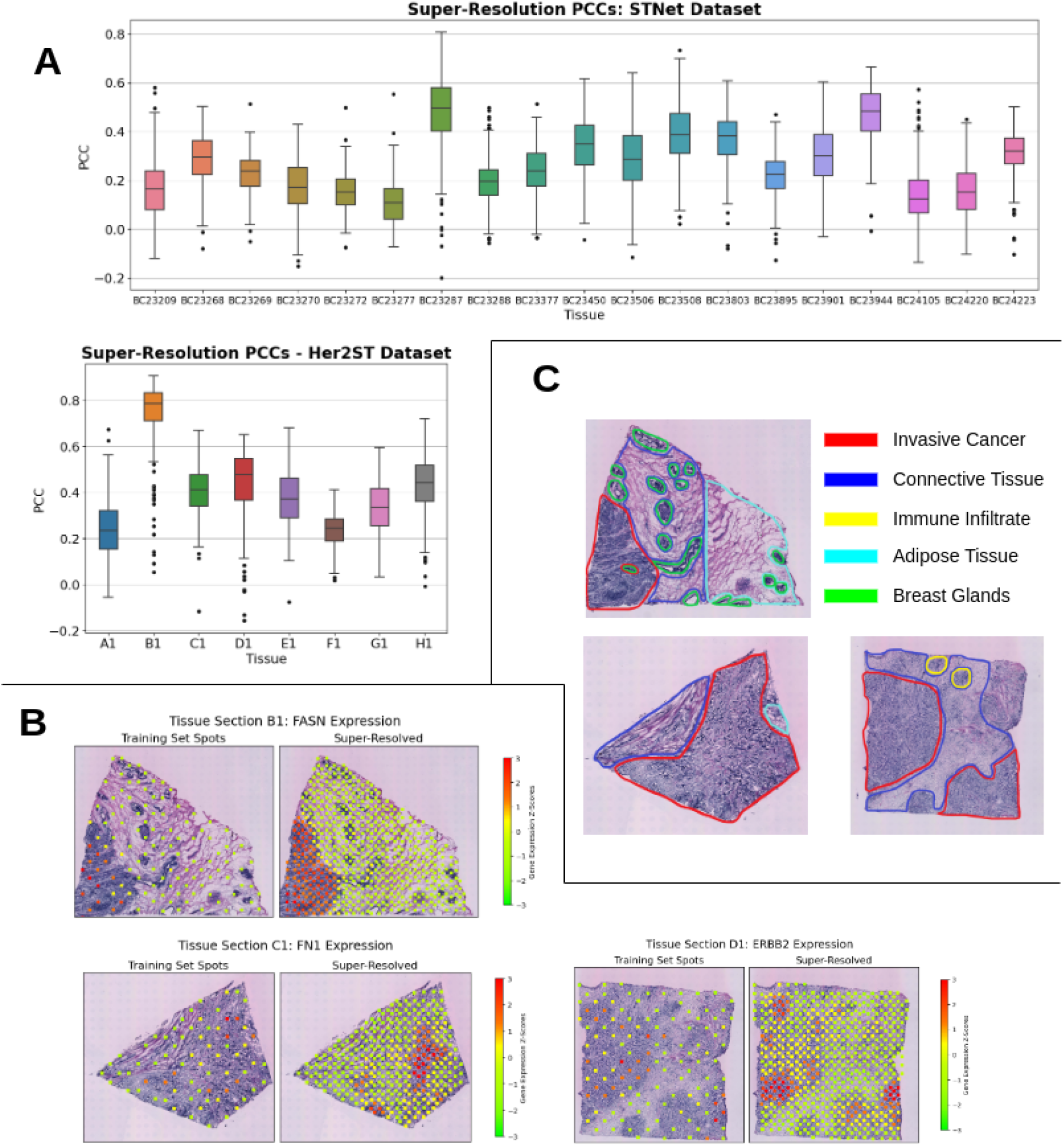
(a) Pearson correlation coefficients between true and STING-inferred gene expression at the spots hidden during training, (b) super-resolved gene expression profiles of the genes **FASN, ERBB2**and **FN1**in tissue sections *B1, C1* and *D1* respectively, from the Her2-ST dataset and (c) pathologist’s annotation of the tissue sections *B1, C1* and *D1*.

We further visualize the training and super-resolved gene expression profiles for **ERBB2, FASN**, and **FN1** (Figure 2b), which are highly associated with breast cancer. **FASN** encodes fatty acid synthase, a pro-oncogenic enzyme commonly overexpressed in various cancers to support increased lipogenesis for sustained proliferation^37^. **FN1** encodes fibronectin-1, a glycoprotein linked to tumor cell migration and metastasis, often overexpressed in pre-metastatic conditions^38^. **FN1** is also implicated in aspartate metabolism for enhanced nucleotide synthesis^39^ and signaling pathways that sustain the tumor microenvironment^40^. **ERBB2** is a key driver of proliferation in Her2-positive breast cancer^41^.

In tissue section *B1*, **FASN** overexpression is observed in regions marked as invasive cancer, with the super-resolved profile closely matching the malignant area. Additionally, clusters of **FN1** and **ERBB2** overexpression align with malignant regions in tissue sections *C1* and *D1*, respectively.

### Tissue-Wide Gene Expression Inference

We infer spatial gene expression across entire tissue sections that STING has not encountered during training. To eliminate potential biases arising from similarities in tissue morphology and gene expression within tissues derived from the same patient, we conduct patient-wise cross-validation. In each training iteration, we exclude all tissues from a specific patient and evaluate the trained model on the withheld tissues.

We benchmark STING against two previously published methods for spatial gene expression inference: `ST-Net` by He et al.^26^ and `TCGN` by Xiao et al.^35^. `ST-Net` employs a purely CNN-based architecture, whereas `TCGN` integrates CNNs, GNNs, vision transformers, and attention mechanisms. While training `ST-Net`, we use a 10× higher learning rate than that used by He et al. to counter slow convergence, ensuring that this adjustment does not lead to instability. For `TCGN`, we reduce the number of training epochs due to computational constraints but still achieve complete loss convergence. We evaluate all three methods on the expression of the 250 most highly expressed genes across both datasets.

The PCCs between true and predicted expression for tissue sections *A1–H1* from the Her2-ST dataset are visualized in Figure 3a. STING consistently outperforms `TCGN` and either surpasses or performs comparably to `ST-Net`. The performance difference is most striking in tissue section *B1*, where STING achieves a mean PCC of 0.69, significantly higher than `ST-Net` (0.45) and `TCGN` (−0.07). Similar trends are observed in sections *C1* (0.34, 0.24, and 0.16 for STING, `ST-Net`, and `TCGN`, respectively), *D1* (0.32, 0.19, and 0.07), and *G1* (0.23, 0.17, and 0.07). While STING performs worse on *E1* and *F1*, `ST-Net` and `TCGN` also yield poor correlations with true expression in these sections.

**Figure 3.**
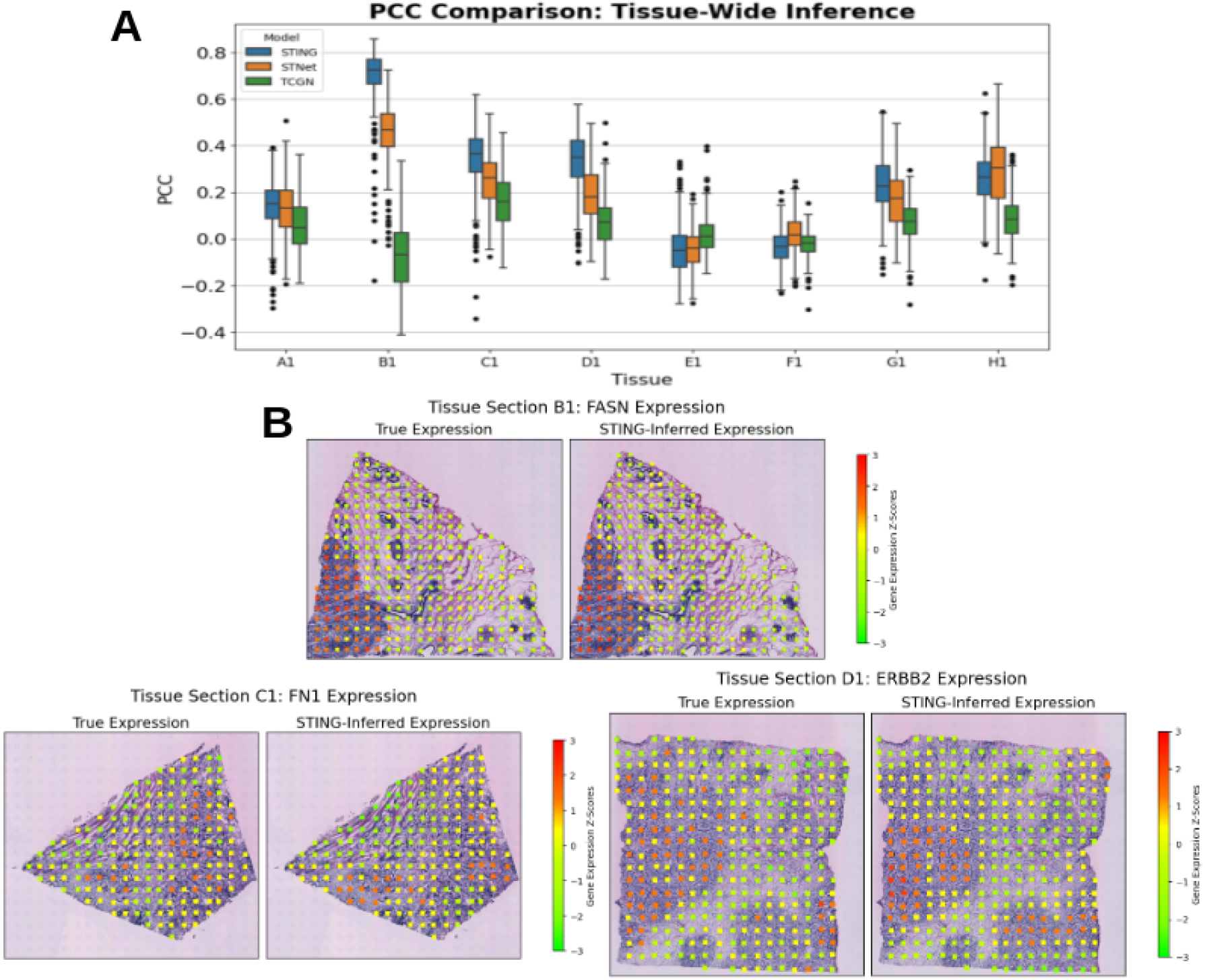
(a) Comparison of Pearson correlation coefficients for gene expression inferred using STING, `ST-Net` and `TCGN`, (b) STING-inferred gene expression for the genes **FASN, FN1** and **ERBB2** in tissue sections *B1, C1* and *D1* respectively

Beyond accuracy, STING offers significant computational advantages, requiring **less time and memory** than both `ST-Net` and `TCGN`. It can be trained and deployed on a standard computer and achieves fast convergence even without a GPU. Table 1 summarizes the mean training times for all three approaches during cross-validation.

**Table 1.** Comparison of mean training times during cross-validation for STING, ST-Net and TCGN.

| Method | Hardware | Number of Training Epochs | Mean Total Training Time | Mean Training Time Per Epoch |
| --- | --- | --- | --- | --- |
| STING | Intel Xeon CPU (2.2 GHz)<br>30 GB Memory | 70 | 683 sec | 9.7 sec |
| ST-Net | NVIDIA A100 GPU<br>30 GB Memory<br>10 GB GPU Memory | 50 | 3613 sec | 72 sec |
| TCGN | NVIDIA A100 GPU<br>60 GB Memory<br>20 GB GPU Memory | 20 | 33124 sec | 552 sec |

The PCCs for all tissue sections across the ST-Net and Her2-ST datasets are provided in Supplementary Figures 3 and 4, respectively. We also visualize the inferred and true expression of **FASN, FN1**, and **ERBB2** in *B1, C1*, and *D1*, respectively (Figure 3b). The predicted expression of **FASN** in *B1* and **ERBB2** in *D1* correlates strongly with observed values, with clusters of overexpression aligning with tumor regions identified by pathologists (Figure 2c). Notably, in *D1*, STING identifies a larger **ERBB2** overexpression cluster in the bottom-right region of the tissue than what is captured by Visium. This overexpression aligns more closely with the pathologist’s annotations, suggesting that STING may help identify errors in Visium-based spatial gene expression measurements, which could arise from low RNA detection efficiency or other limitations.

Another key advantage of STING is its ability to generate smooth, biologically consistent spatial gene expression profiles. Unlike alternative methods, STING-inferred expression maps are **free from abrupt changes**, aligning with the biological principle that spatially adjacent cells exhibit similar gene expression patterns.

To further assess STING’s generalization capability, we train STING on the ST-Net dataset and evaluate it on the Her2-ST dataset, and vice versa. The resulting PCCs for *A1–H1* from the Her2-ST dataset are shown in Figure 4, with results for all tissues provided in Supplementary Figures 5 and 6. The PCC distributions closely mirror those observed in patient-wise cross-validation, reinforcing that STING generalizes well across diverse breast cancer subtypes and is not limited to a specific dataset.

**Figure 4.**
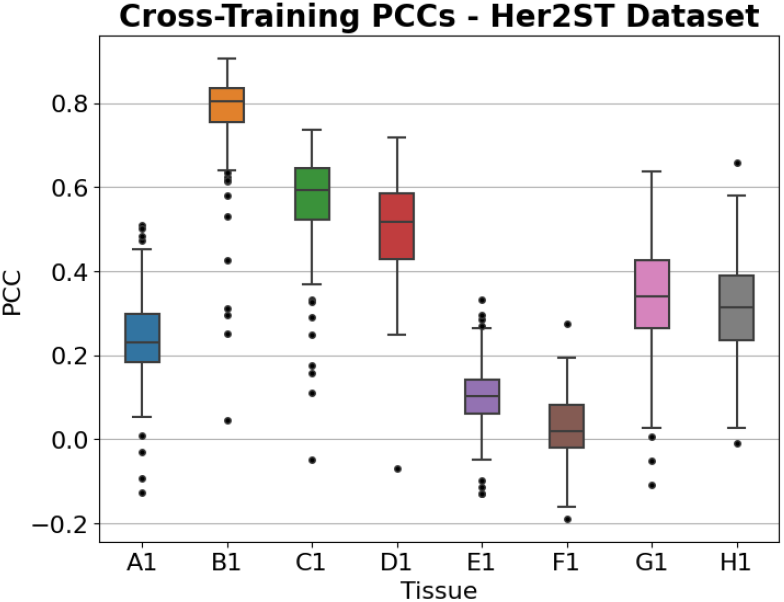
Pearson correlation coefficients for predicted gene expression in tissues *A1-H1* after training STING on tissues from the ST-Net dataset

## Discussion

STING predicts spatial gene expression by leveraging morphological features of cells and their neighbors, extracted from readily available histopathology images. In our experiments, STING successfully super-resolves sparse spatial gene expression profiles, predicting expression at up to five times the number of spatial locations used for training. The super-resolved profiles align well with held-out spots, achieving a mean PCC above 0.35 for six out of eight tissues in the Her2-ST dataset. Furthermore, STING generalizes effectively across entire tissue sections withheld during training, both within and across datasets, demonstrating robustness to variations in sample preparation, tissue staining, and experimental protocols. Importantly, STING-predicted spatial gene expression patterns correlate well with pathologists’ annotations of breast cancer tissues. Although our experiments focus on breast cancer datasets, we expect STING to generalize to other cancer types and non-cancerous tissues, provided high-resolution histopathological images are available.

Predicting molecular properties of cancer such as mutation status or gene expression has been a topic of great interest. At a molecular level, cancer is inherently heterogeneous, with tumors consisting of diverse cell types and regions exhibiting varying gene expression patterns. SRT enables oncologists to map gene expression across tumor regions, uncovering molecular details that aid in clinical decision-making and treatment planning. Models like STING facilitate the inference of cancer-related gene expression, potentially contributing to better patient outcomes in oncology.

The tumor microenvironment plays a crucial role in cancer progression, treatment response, and drug resistance. Deciphering its molecular composition is essential for understanding mechanisms of metastasis and immunotherapeutic resistance, where computational approaches like STING for spatial gene expression inference can be highly beneficial.

Beyond oncology, spatial transcriptomics complements single-cell RNA sequencing (scRNA-seq) in providing deeper insights into RNA expression mechanisms. While scRNA-seq offers high transcriptomic resolution at the single-cell level, it lacks spatial context. SRT maps transcript locations in tissues, enriching scRNA-seq data by enabling better cell type mapping and spatial organization analysis. Conversely, scRNA-seq can fill gaps in SRT data, such as missing gene expression information, making these two approaches highly synergistic.

As the availability of spatial transcriptomic data as well as other spatial omics data grows, so do the modeling opportunities. Wang et al.^42^ in their review on deep learning-driven inference of spatial omics data state that methods capturing both local and global features do better than ones simply looking at the local neighborhood of spots, which is consistent with our findings. Additionally, models imputing spatial gene expression of additional genes from the expression profiles of a smaller set of genes^43,44^ further increase the value of approaches like STING, which are limited to only a small subset of genes due to the lack of signal-to-noise ratio. Further advancements in STING-like models depend on access to additional spatial transcriptomic datasets and standardization of experimental methodologies, including tissue staining and imaging procedures. By computationally inferring spatial gene expression directly from histopathological images, STING reduces reliance on multiple invasive procedures and expands clinical possibilities for non-invasive molecular profiling, paving the way for more precise and accessible diagnostic and prognostic tools.

## Methods

### Data Preprocessing

We remove 12 tissues from 4 patients in the ST-Net dataset due to their overlap with the Her2-ST dataset. Additionally, many genes in both datasets exhibit low or undetected expression, likely due to RNA detection errors in the 10x Visium platform. These low-expression genes could introduce noise and mislead the model during training. To maximize the signal-to-noise ratio, we select 250 genes with the highest mean expression independently for each dataset.

Given the wide range of gene expression values, we apply a logarithmic transformation after adding a pseudo-count of 1 to account for genes with zero expression. We do not apply any additional scaling, ensuring that predicted gene expression values can be easily scaled back without additional parameters.

### STING Design

We employ a GNN model to infer gene expression at various spatial locations within tissues. The model is trained using gene expression data and corresponding tissue WSIs, which are represented as graphs to capture spatial relationships.

#### Graph Data

Each tissue in the dataset is represented as a graph consisting of nodes and edges. The spatial locations where gene expression is measured, referred to as “spots”, form the nodes of the graph (see Figure 1a). Each node is connected to “*k*” nearest neighbors based on Euclidean distance, defining the edges (see Figure 1a). The edge weights correspond to the distance between connected spots (see Figure 1b).

To incorporate morphological features, we extract a *W × H* pixel area around each spatial location in the WSI where gene expression is measured. These extracted regions are processed through `KimiaNet`^45^, a `DenseNet-121`-based CNN pre-trained on WSIs. We download `KimiaNet`’s pre-trained weights and apply them to `DenseNet-121`, removing its fully connected layer (see Figure 1c). The WxH sub-image is then passed through `KimiaNet` to generate 1024-dimensional embeddings, which serve as node features (see Figure 1).

Thus, each graph representation of a tissue consists of nodes (spots), node features (CNN embeddings), edges (spatial proximity), and edge weights (Euclidean distance).

#### GNN Model

We employ a GNN model with two sequential graph convolutional layers^46^ to predict spatial gene expression. The tissue graphs serve as input to the first graph convolutional layer, which transforms the 1024-dimensional node features into a 256-dimensional latent space, followed by a rectified linear unit (ReLU) activation. The second graph convolutional layer further processes the latent space to output the predicted gene expression scores. To prevent overfitting, we apply dropout to the first graph convolutional layer. Figure 5 illustrates the architecture of the GNN model.

**Figure 5.**
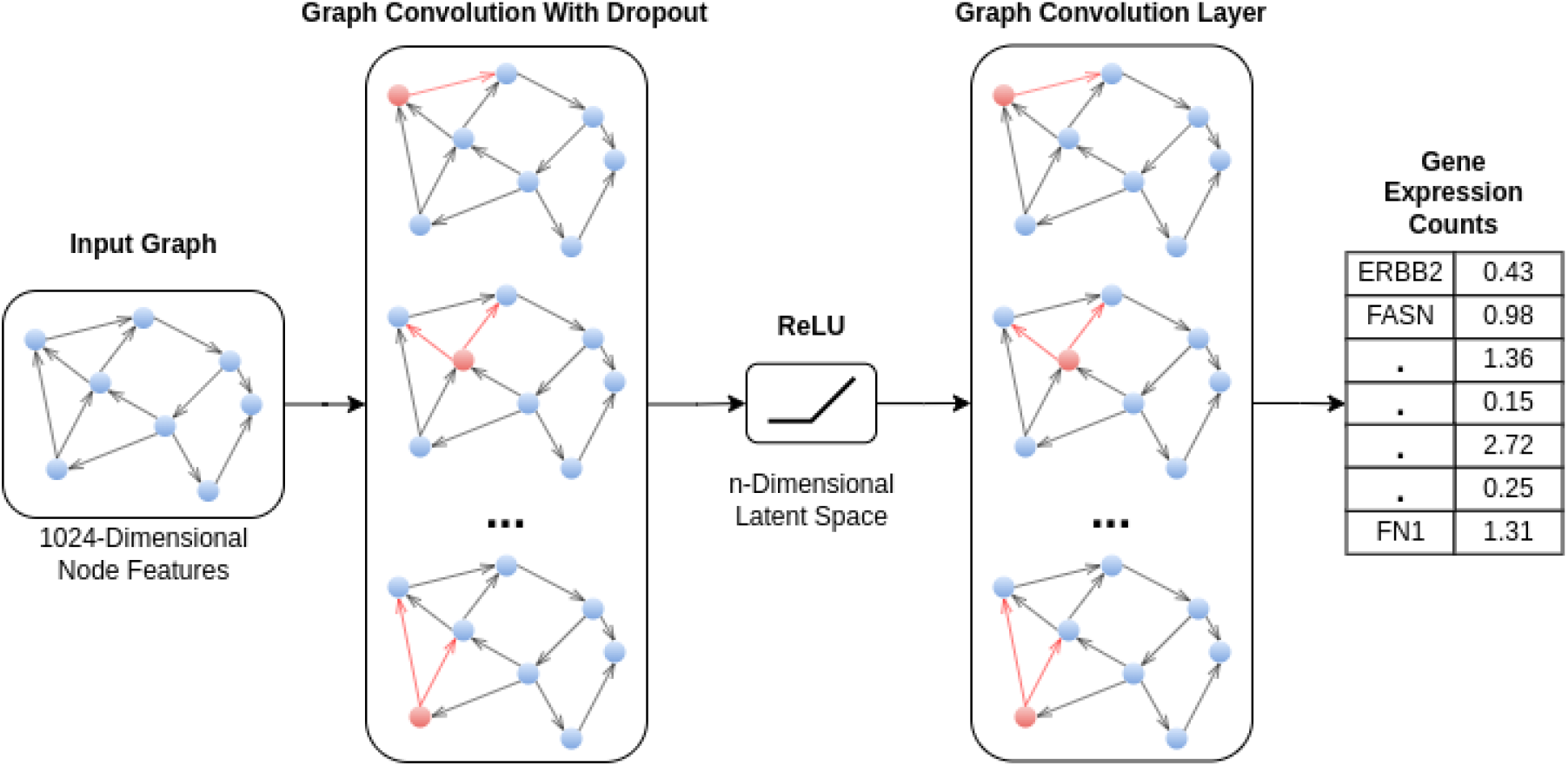
Architecture of the GNN model. The input graph corresponding to a tissue is passed through a graph convolutional layer, which converts the 1024-dimensional node features (generated using `KimiaNet`) to a n-dimensional latent space. The latent space serves as an input to the second graph convolutional layer, which outputs predicted gene expression values.

### Super-Resolving Gene Expression

We leverage STING to super-resolve gene expression within a tissue. During training, STING operates on a graph constructed using 40% of the available spots in a tissue, selected via farthest point sampling to ensure optimal tissue coverage. After training, we recompute the graph, incorporating the previously excluded spots, and use STING to predict gene expression scores.

To assess super-resolution accuracy, we compute Pearson correlation coefficient (PCC) and mean squared error (MSE) between the true and predicted gene expression scores at the previously hidden spots. Additionally, we sample extra spots, totaling up to five times the number of training spots, and visualize the super-resolved gene expression profiles.

For super-resolution, we train the graph convolutional network (GCN) for 200 epochs using the Adam optimizer with a learning rate of 0.001, apply a weight decay of 0.0005, and minimize the Huber loss function, expressed as:

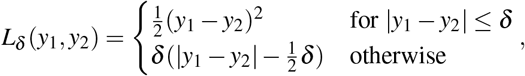

where *y*_1_ and *y*_2_ are the true and predicted gene expression values, respectively, and *δ* is a threshold parameter that determines the transition between quadratic and linear loss regions. For our experiments, we set *δ* = 1, its default value.

This loss function is robust to outliers, making it well-suited for gene expression prediction, where erroneous or noisy measurements can occur due to experimental variability.

### Predicting Tissue-Wide Gene Expression

We assess STING’s ability to infer spatial gene expression in tissues not encountered during training. First, we perform patient-wise cross-validation on the Her2-ST dataset, systematically excluding all tissues from one patient in each iteration while training the model on the remaining samples. The trained model is then evaluated on the excluded tissues, and this process is repeated for all eight patients in the dataset. Next, to ensure that STING’s predictive performance is not confined to a specific dataset, we train the model on all tissues in the Her2-ST dataset and evaluate its performance on the ST-Net dataset.

## Data Availability

The raw sequencing files for the Her2-ST data are available with restricted access at the European Genome-Phenome Archive (EGA) with the identifier EGAD00001008031, and access can be obtained by contacting the authors of the original publication^36^. The processed count matrices and corresponding H&E images are available at https://doi.org/10.5281/zenodo.4751624.

The ST-Net data are available at https://data.mendeley.com/datasets/29ntw7sh4r/5.

## Author contributions statement

M.B. and K.K. outlined the study. K.K. carried out the experiments and analyses with insights from M.B. and A.R., and M.B., K.K. and A.R. drafted the final manuscript.

