## Supplementary Figures for "STING: A Graph Neural Network Approach for Computational Inference of Spatial Transcriptomic Profiles"

**(Supplementary Information)**

Kaushik Karambelkar	Arvind Rao	Mayank Baranwal

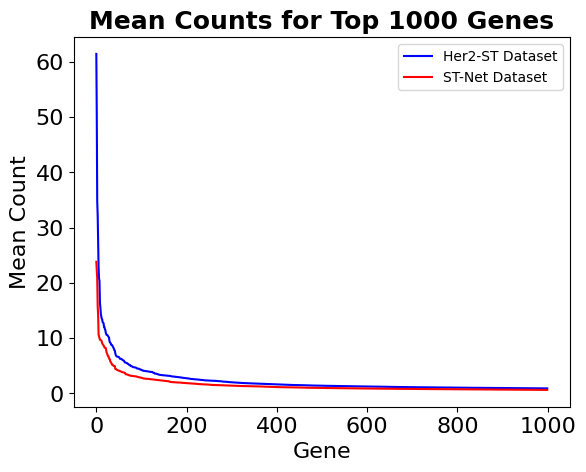

**Supplementary Figure 1**: Mean transcript counts for the top 1000 genes in both datasets

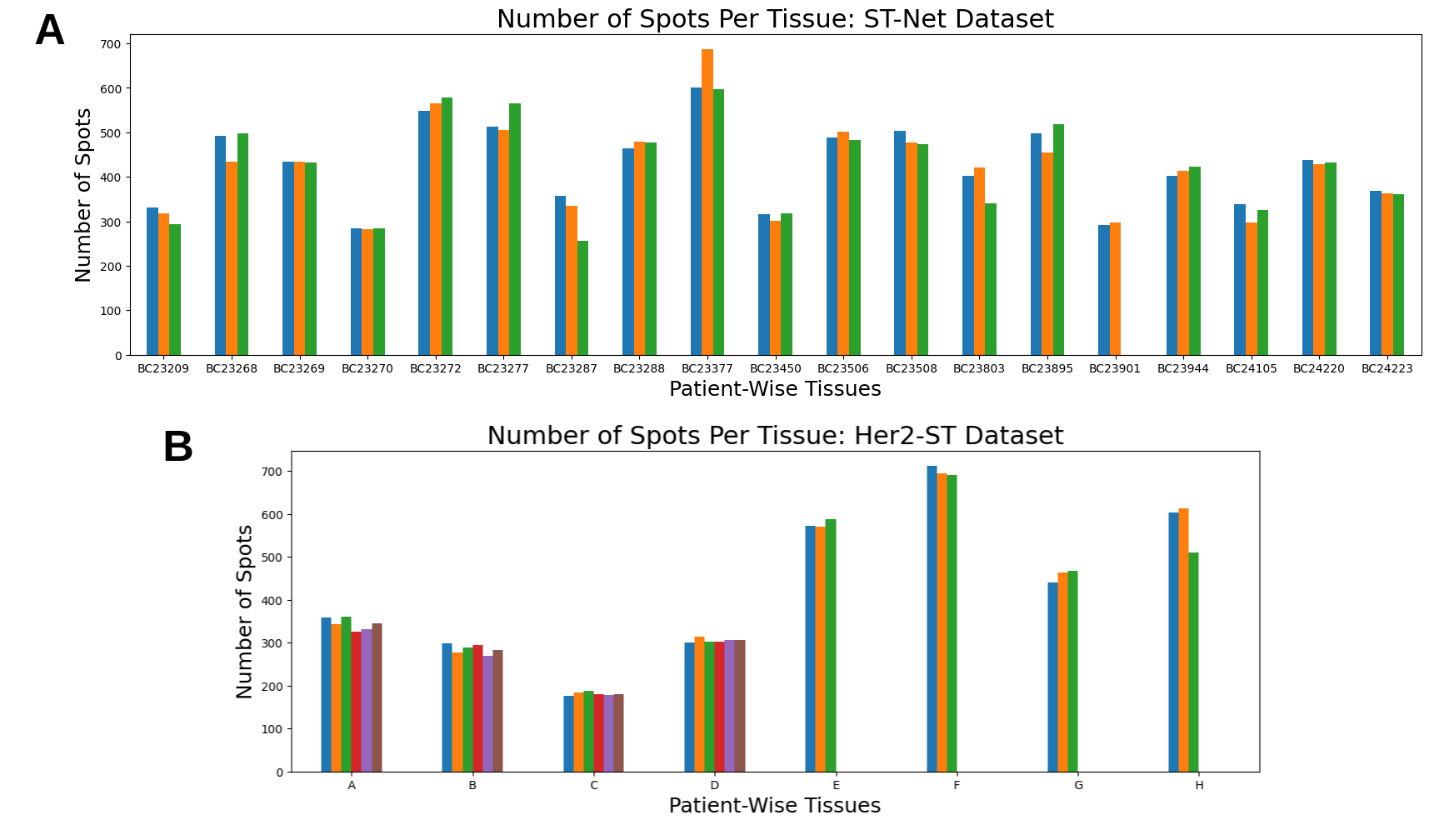

**Supplementary Figure 2**: Number of spots with gene expression measurements in (A) the ST-Net dataset and (B) the Her2-ST dataset.

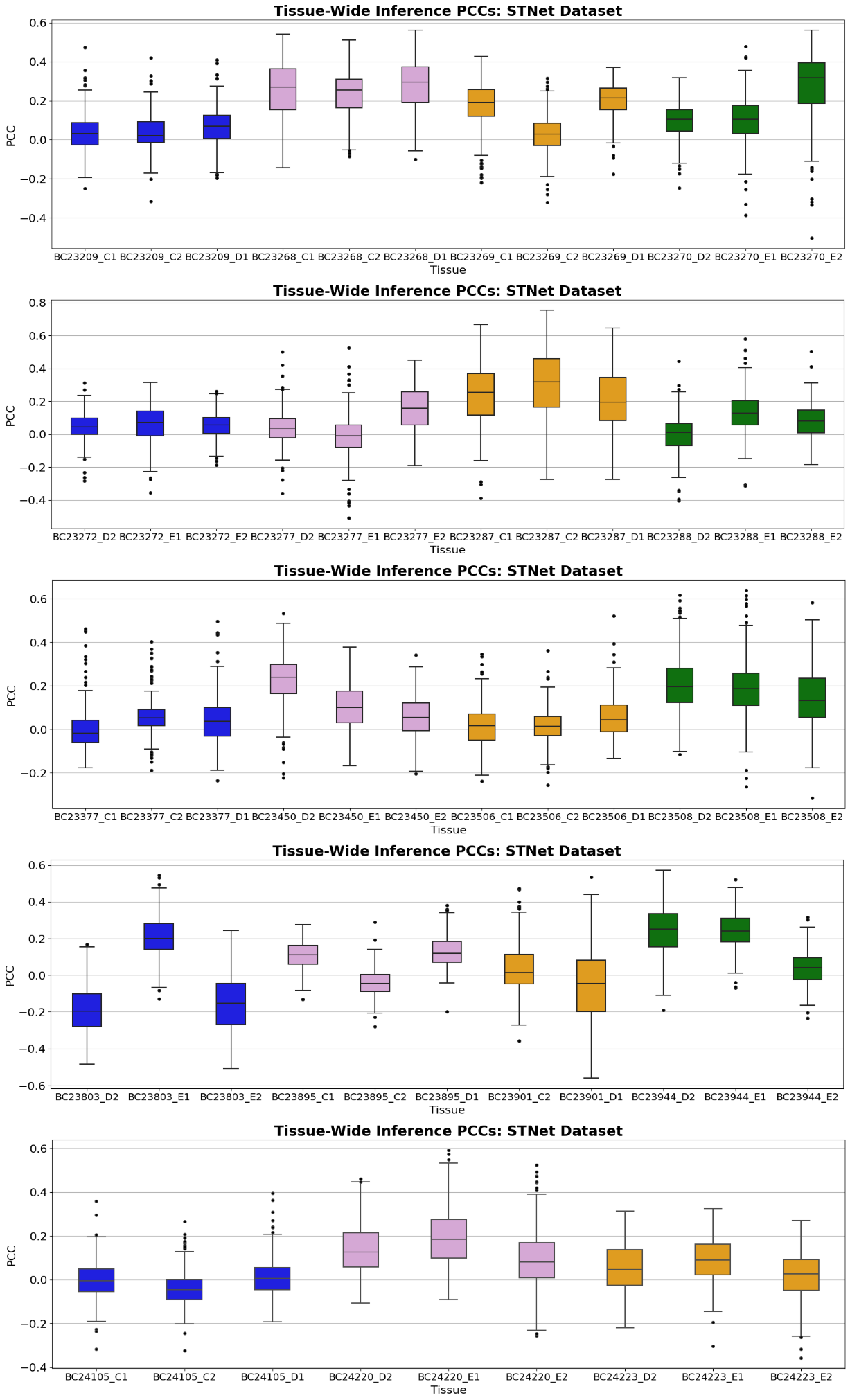

**Supplementary Figure 3**: Box plots of the mean Pearson correlation coefficients between true and predicted gene expression for the 250 genes in each tissue in the ST-Net dataset

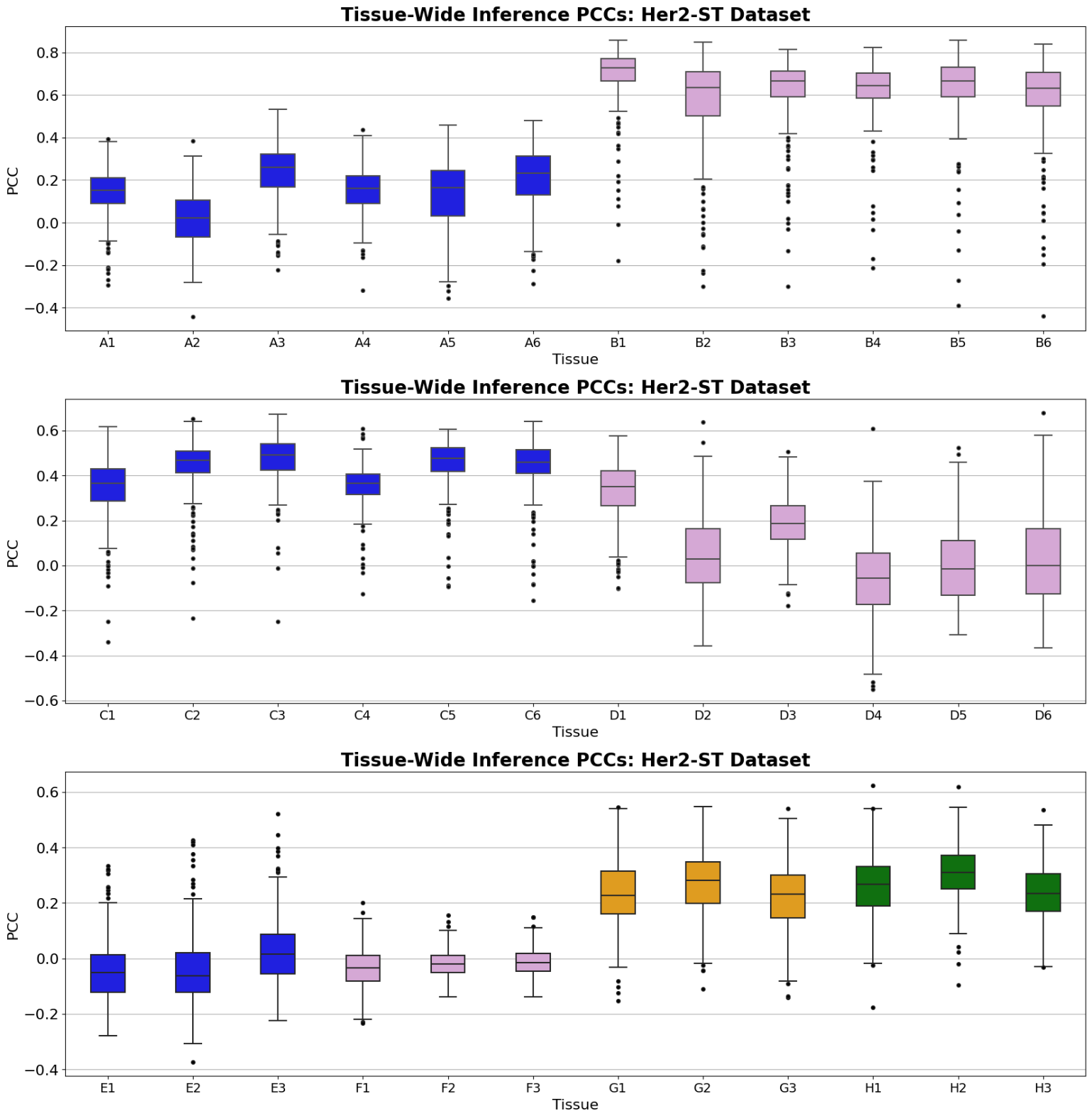

**Supplementary Figure 4**: Box plots of the mean Pearson correlation coefficients between true and predicted gene expression for the 250 genes in each tissue in the Her2-ST dataset

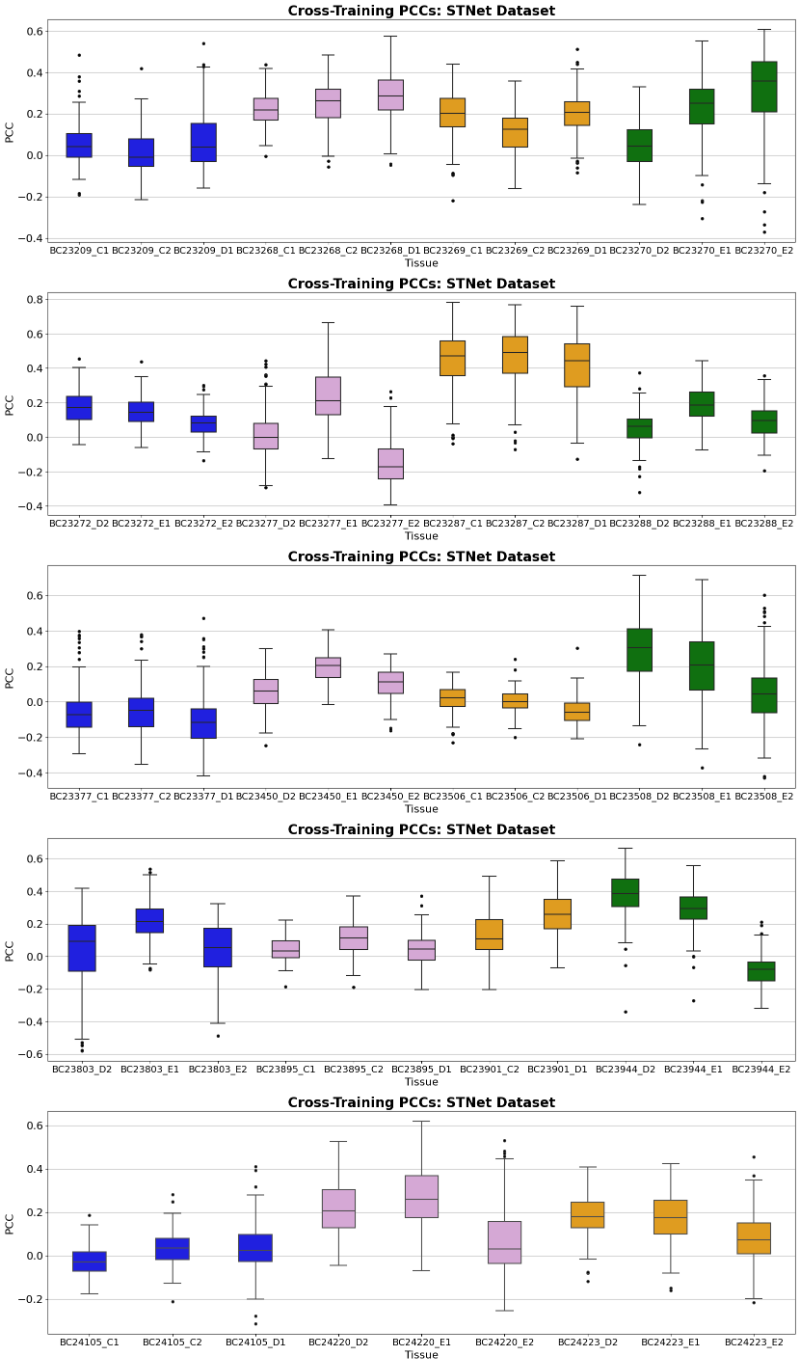

**Supplementary Figure 5**: Box plots of the mean Pearson correlation coefficients between true and predicted gene expression for the 250 genes in each tissue in the ST-Net dataset after training STING on the Her2-ST dataset.

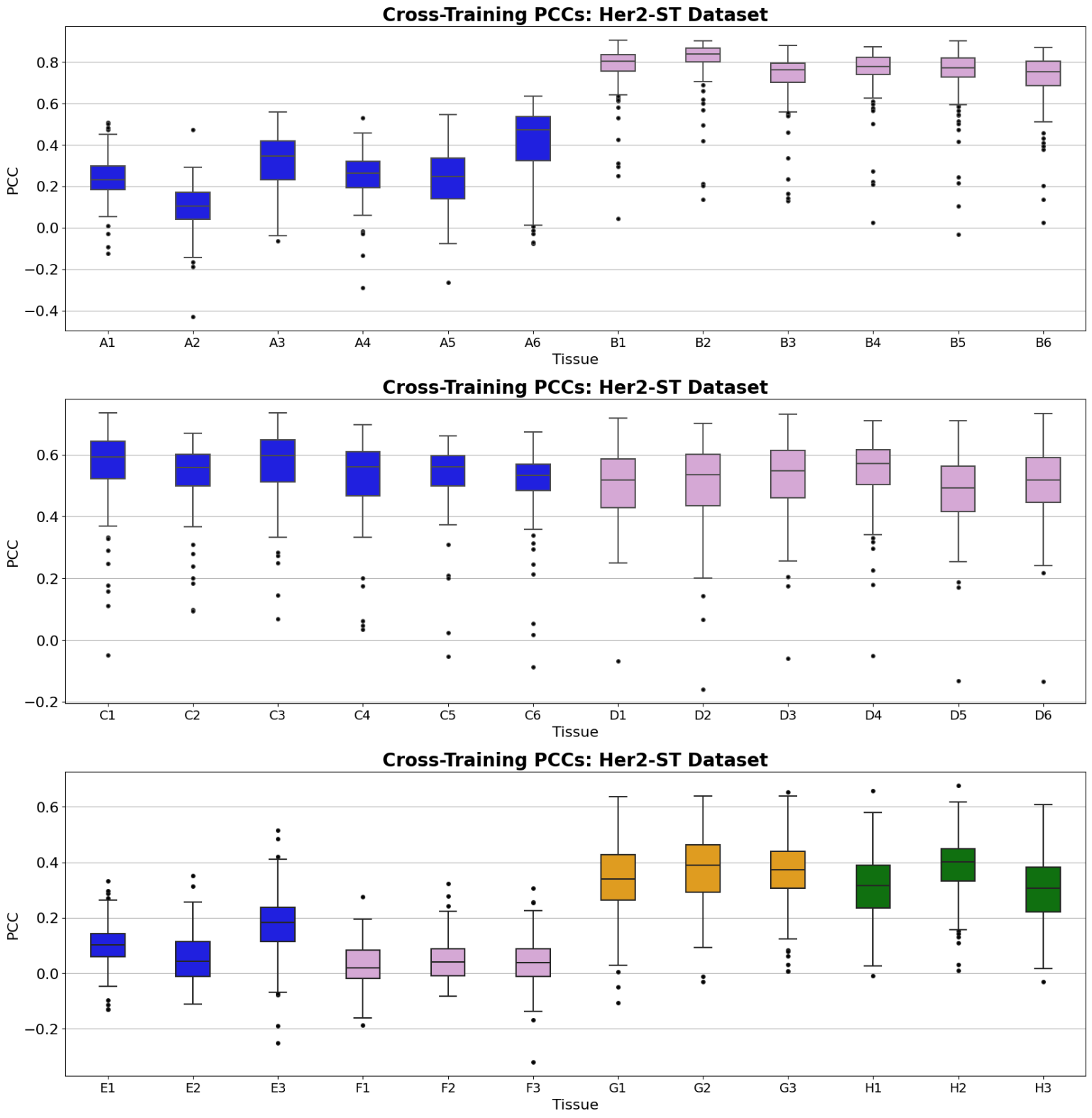

**Supplementary Figure 6**: Box plots of the mean Pearson correlation coefficients between true and predicted gene expression for the 250 genes in each tissue in the Her2-ST dataset after training STING on the ST-Net dataset.
